# Effect of adiposity on leukocyte telomere length in US adults by race/ethnicity: The National Health and Nutrition Examination Survey

**DOI:** 10.1101/661819

**Authors:** Sharon K. Davis, Ruihua Xu, Rumana J. Khan, Amadou Gaye, Yie Liu

## Abstract

**Objective:** Obesity is associated with telomere attrition – a marker of cellular and biological aging. The US has the highest proportion of obesity and is comprised of a racially/ethnic diverse population. Little is known about the relationship between obesity and telomere attrition according to race/ethnicity in the US. Our objective is to examine the differential association.

**Design and setting:** The effect of body mass index (BMI), % total body fat (TBF) and waist circumference (WC) on leukocyte telomere length (LTL) were examined as adiposity measures according to race/ethnicity and sex specific race/ethnicity using separate adjusted linear regressions on a sample of 4,919 respondents aged 20-84 years from cross-sectional 1999-2002 data using the US National Health and Nutrition Examination Survey. Mediation analyses assessed health behaviors associated with relationship between adiposity measures and LTL.

**Main outcome measure:** LTL

**Results:** African Americans (AA) experienced a 28% and 11% decrease in LTL associated with increasing BMI and WC, (*p*=.02 and .03) respectively. Mexican Americans (MA) experienced a 33% decrease in LTL associated with increasing %TBF (*p*=.04). Whites experienced a 19%, 23%, and .08% decrease in LTL associated with increasing BMI, %TBF, and WC, (*p*=.05, .003, .02) respectively. White men experienced a 26% decrease in LTL due to increasing BMI (*p*=.05). AA women experienced a 41%, 44%, and 16% decrease in LTL due to increasing BMI, %TBF, and WC, respectively (*p*=.007, .02, .04). White women experienced a 29% decrease in LTL associated with increasing %TBF (*p*=.006). Selected health behaviors were associated with the relationship between adiposity measures and LTL.

**Conclusion:** Overall, AA and Whites have worse cellular and biological aging related to collective adiposity measures. According to sex, AA women experienced more deleterious cellular and biological aging. Findings suggest tailored interventions to improve adverse behaviors that contribute to obesity may improve telomere attrition in US adults.

## Introduction

Secular rates of risk factors associated with cardiovascular disease (CVD) have been declining among US adults across all racial/ethnic groups.[1] Obesity rates, on the other hand, have increased. [2] Worldwide obesity has nearly tripled since 1975.[3] The US has the highest proportion of obesity.[2] More than one-third of US adults are obese.[4] Prevalence rates differ by race/ethnicity and by sex according to race/ethnicity.[4] Obesity is a major risk factor for many age-related CVD chronic conditions such as hypertension, type 2 diabetes and dyslipidemia which increases the risk for heart failure, heart attack and stroke.[5] It is the leading cause of preventable deaths globally and occurs, in part, due to adverse modifiable lifestyle behaviors such as sedentary physical activity and unhealthy diet.[3]

Telomeres are the DNA-protein complex at the ends of chromosomes.[6] It consists of highly conserved tandem hexameric nucleotide repeats (TTAGGG). Telomere are needed for the replication of DNA and provides protection to chromosomes from nuclease degradation and cellular senescence which promotions the integrity and stability of chromosomes. During the cellular process, telomeres progressively shorten with each cell division. When telomeres shorten to a critical length, replicative senescence is triggered resulting in cell-cycle arrest.[7] In human peripheral leukocytes, telomere shortening has been demonstrated to be a maker for cellular and biologic aging as well as a biomarker for age-related diseases such as CVD.[8] Evidence shows that the pathways through which obesity promotes morbidities include increasing systemic inflammation and oxidative stress; inflammation and oxidative stress have also been linked to telomere attrition.[9-11] Studies have investigated the relationship between adiposity and telomere length. Such studies have produced equivocal results [12-20]. Studies have also examined the association between adverse lifestyle health behaviors and telomere length with similar mixed results. [19, 21-26] One factor that may account for these conflicting findings may be inadequate statistical power due to small sample size, study design, and sample characteristics. Little is known about racial/ethnic differences between adiposity and telomere length. In addition, we are unaware of any studies that have investigated adverse health behaviors as a pathway associated with adiposity and telomere length. The objective of our research was to examine a large US representative, socioeconomically and racially/ethnic diverse population. This represents the first study to examine adiposity and telomere length according to race/ethnicity, sex according to race/ethnicity and mediating pathways due to adverse lifestyle in the US. We hypothesize that the association between telomere length and adiposity will be moderated by race/ethnicity and sex. We further hypothesize that adverse health behaviors will have a mediating effect between adiposity and telomere length. Findings will provide important information about the rate of biological aging due to telomere length and adiposity across major race/ethnic groups in the US and according to corresponding sex; as well as the implications associated with adverse health behaviors.

## Materials and Methods

### Study design and sampling procedures

Data was collected from the 1999-2000 and 2001-2002 cycles of the National Health and Nutrition Examination Survey (NHANES). This is a nationally representative cross-sectional survey and physical examination of civilian, noninstitutionalized US population conducted by the US Centers for Disease Control and Prevention (CDC) since 1960.[27] NHANES utilizes a 4-stage sampling design which includes 1) primary sampling units (PSUs) consisting of single counties, 2) area segments within PSUs, 3) households within segment areas, and 4) persons within households. On average 2-3 individuals per household were sampled. NHANES 1999-2002 oversampled low-income individuals, African Americans and Mexican Americans to obtain more accurate estimates in these populations. All respondents aged ≥20 during this period were asked to provided DNA specimens to establish a national probability sample of genetic material for future research. DNA from the most recent NHANES is only available in the form of crude lysates of cell lines thereby precluding the assay of leukocyte telomere length (LTL). However, DNA collected during 1999-2002 is purified from whole blood thus facilitating the assay of LTL. Pooled data were available for public download (http://www.cdc.gov/nchs/nhanes_questionnaires.htm).

Of the 10,291 respondents eligible to provided DNA, 7,825 provided DNA and consented to future genetic research. We excluded 653 respondents whose self-reported race/ethnicity was “other” or “other Hispanic,” since our goal was to examine more discrete self-reported race/ethnic groups (i.e. White, African American, Mexican American). We also excluded 225 respondents aged ≥85 because of survival bias among the extreme elderly.[28] An additional 2,037 were excluded due to missing data on one or more variables in the models - resulting in a final sample size of 4,919. There were no significant sociodemographic differences between the full sample and the final sample. Sampling weights were used to address oversampling and non-response bias and to ensure that estimates are representative of the general US population. Written informed consent was obtained from each participant. Human subject approval was provided by the Institutional Review Board (IRB) at the CDC and the study protocol was approved by the IRB of the National Institutes of Health.

### Data and data collection

Aliquots of purified DNA were provided by the laboratory of the CDC. DNA was isolated from whole blood using the Puregene (D-50K) kit protocol and stored at −80°. The LTL assay was performed in the laboratory of Dr. Elizabeth Blackburn at the University of California, San Francisco, using the quantitative polymerase chain reaction (PCR) method to measure telomere length relative to standard reference DNA (T/S ratio).[29] The single-copy gene was used as a control to normalize input DNA was human beta-globin. Each sample was assayed twice. T/S ratios that fell into the 7% variability range were accepted; the average of the two was taken as the final value. A third assay was run for samples with greater than 7% variability and the average of the two closest T/S values was used. The inter-assay coefficient of variation was 4.4%.

Body mass index (BMI), estimated % total body fat, and waist circumference were analyzed separately as measures of adiposity. BMI was calculated as weight in kilograms divided by height in meters squared (kg/m^2^) using a calibrated electronic digital scale and a stadiometer. Estimated % total body fat was assessed using duel-energy X-ray absorptiometry of the whole body that lasted 3 minutes (Hologic scanner, QDR-4500, Bedford, MA, USA). Total % body fat was calculated as total body fat mass divided by total mass × 100. Waist circumference was measured in centimeters using a tape measure around the trunk, at the iliac crest, crossing at the mid-axillary line. The details of these assays have been described elsewhere.[30]

Adverse health behaviors were assessed as pathway mediators between adiposity exposures and LTL outcome; these include smoking, drinking, physical activity, and diet. Smoking was measured a cumulative exposure to tobacco smoke in pack-years, calculated as the average number of cigarettes smoked per day times the number of years smoked divided by 20 (number of cigarettes in one pack).[31] Dummy variables were 30-=>59 pack years, <30 pack years, and never smoked was coded as the reference. Drinking was based on daily alcohol consumption defined as heavy, moderate and abstainer.[27] Heavy drinkers were defined as women reporting having drunk ≥2 alcoholic beverages in the past 12 months per day and men reporting having drunk ≥3 alcoholic beverages in the past 12 months per day. Moderate drinkers were defined as women reporting <2 drinks per day in the past 12 months and men reporting <3 drinks per day in the past 12 months. Men and women reporting no alcoholic beverages in the past 12 months per day were the reference and defined as abstainers. Physical activity was based on guidelines provided by the Department of Health and Human Services.[32] Respondents met or exceeded recommended guidelines if they reported ≥150-≥300 minutes per week of physical activity, such as brisk walking, gardening, and muscle-strengthening based on total number of minutes reported for each activity. Those reporting <150 minutes of physical activity per week were below the recommended guidelines. Diet was based on The Healthy Eating Index (HEI) developed by the US Department of Agriculture in 2005.[33] The score is the sum of 10 components representing different aspects of a healthy diet. Each component of the index has a maximum score of 10 and a minimum score of zero. The maximum overall score for the 10 components combined is 100. An overall index score ≥ 80 implies a “good” diet, an index score between ≥51and 80 implies a diet that “needs improvement,” and an index score <51 implies a “poor” diet.

Race/ethnicity was based on self-reported non-Hispanic White, non-Hispanic Black and Mexican American thereto referred to as White, African American and Mexican American. Confounding demographic variables that may affect the relationship between adiposity and telomere length included age in years at the time of the survey, age^2^, sex, socioeconomic status based on Poverty Income Ratio (PIR), adiposity related health outcomes, markers of inflammation and oxidative stress, and characteristics of the blood from which DNA was extracted. PIR was calculated as the ratio of income to the poverty threshold for a household of a given size and composition. PIR values below 1.00 are below the official poverty threshold as defined by the US Census Bureau.[34] Adiposity related health status was based on respondents answer to the questions “have you ever been told by a doctor or other health professional that you had hypertension, also called high blood pressure and “have you ever been told by a doctor or health professional that you have diabetes or sugar diabetes?” Markers of inflammation and oxidative stress included C-reactive protein (CRP) and gamma glutamyltransferase (GGT) measured from serum. Characteristics of the blood samples from which DNA was extracted included white blood cells (µL), lymphocytes (%), monocytes (%), neutrophils (%), eosinophils (%), and basophils (%).

### Statistical methods

Descriptive analysis was performed stratified by total sample, African American, Mexican American and White according to study variables. Continuous variables are presented as means ± standard deviation based on ANOVA and categorical variables as percent based on chi-square. Leukocyte telomere length was log-transformed by natural logarithm prior to modeling. Multivariate linear regression models were fitted to assess the relationship between LTL and each adiposity measure. We report the percentage change in the average value of telomere length for a one-unit change in a predictor variable based on the beta estimate for telomere length as the outcome. All regression models accommodated the complex sampling design of NHANES by incorporating strata and PSU indicators, as well as sample weights for the genetic subsample.[35] We first compared the association between LTL and each adiposity measures stratified by each race/ethnic group. We then stratified separately by sex according to race/ethnic group. The models for stratified race/ethnic groups was adjusted for age, age^2^, sex, PIR, hypertension status, type 2 diabetes status, CRP, GGT, white blood cells, lymphocytes, monocytes, neutrophils, eosinophils, and basophils. Sex, hypertension status and type 2 diabetes status were entered as categorical variables. Age, age^2^, PIR, CRP, GGT and the characteristics of blood was entered as continuous variables. The use of age along with a age^2^ term is important when analyzing LTL given the strength of its association with age and the potential for nonlinearity in this association. The models stratified by men and women according to race/ethnicity included the same adjustments as those for the race/ethnicity models – excluding sex.

To assess the moderating effect of race/ethnicity and sex, we entered an interaction term for each adiposity measure in a total aggregate model. We also examined moderating effects by assessing stratified models separately according to race/ethnicity and sex by comparing corresponding parameter estimate across race/ethnic groups and sex using *z*-test (African American versus White, African American versus Mexican American, White versus Mexican American, men versus women). A significant *z*-test suggest moderating effects of race/ethnicity and sex.[36] The mediating effect of each adverse behavior between adiposity measure and LTL was tested separately with a series of regression models using methodological extensions to accommodate categorical mediators.[37, 38] Models included confounding variables adjusted in the aggregate total model. We calculated Arioan test using standardized coefficients of the indirect effects of adiposity on LTL through smoking, drinking, physical activity, and diet. A significant Arioan *z* test suggests a significant indirect effect of adiposity measure and LTL via a candidate mediator. [38] We calculated the proportion of each of the mediators associated with the individual adiposity measure and LTL.[39] We also fitted all 4 mediators as covariates in the final regression model. The mathematical equation formula for Arioan *z* test is:

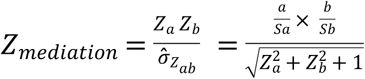

All analyses were conducted using SAS version 9.3.[40] A two-tailed level of significance was established as *P*≤.05.

## Results

Descriptive results in Table 1 reveal African Americans have longer mean LTL (1.13 T/S) compared to the total sample, Mexican Americans, and Whites (1.06, 1.04, 1.05 T/S, respectively). The average mean age for the total sample and Whites was somewhat similar (46 versus 47 years) with younger age observed in African Americans and Mexican Americans (42 versus 38 years). The distribution of women and men was also comparable in the total sample and among Whites (51% versus 49%). African Americans had a higher proportion of women; a lower proportion was in Mexican Americans (53% versus 46%). The prevalence that lived below poverty was lower in the aggregate sample and among Whites (18% and 14%) and a higher prevalence was among African Americans and Mexican Americans (32% and 31%). Mean body mass index and mean waist circumference was slightly higher among African Americans (29kg/m^2^ and 96cm) compared to the other groups; mean % total body fat was similar across all groups. A lower prevalence of African Americans and Mexican Americans smoked 30-≥59 pack years of cigarettes (3.3% and 1.9%); prevalence of non-smokers was also higher in these groups (61% and 62%). The prevalence of all groups smoking <30 pack years of cigarettes was somewhat similar. A lower prevalence of African Americans were heavy drinkers (18%) compared to a higher prevalence among Mexican Americans (33%). The prevalence of moderate drinking was higher in the total sample and among Whites (50% and 53%). The prevalence of abstainers was higher among African Americans (44%). A higher prevalence of African Americans and Mexican Americans were below the recommended level of physical activity per week (56% and 53%). The aggregate sample and Whites had a higher prevalence that met/exceeded the recommended guidelines (61% and 65%). The mean HEI score was somewhat comparable across groups, except slightly lower in African Americans. However, the prevalence of hypertension was higher among African Americans and lower among Mexican Americans (33% versus15%) but comparable in the total sample and in Whites (25%). Mexican Americans as well as African Americans had a higher prevalence of type 2 diabetes (7% and 8%) compared to a comparable lower prevalence in Whites and the total sample (5%). Mean CRP and GGT was higher among African Americans (0.51mg/dL and 42U/L). White blood cell count was lower in African Americans and higher in Mexican Americans compared to the other groups (6.4µL versus 7.30µL). The % of lymphocytes, monocytes, and basophils was higher in African Americans (35%, 8.5%, .68%, respectively). However, percentages of neutrophils and eosinophils were higher among Whites and the total sample (59.3% and 2.7%).

**Table 1:**
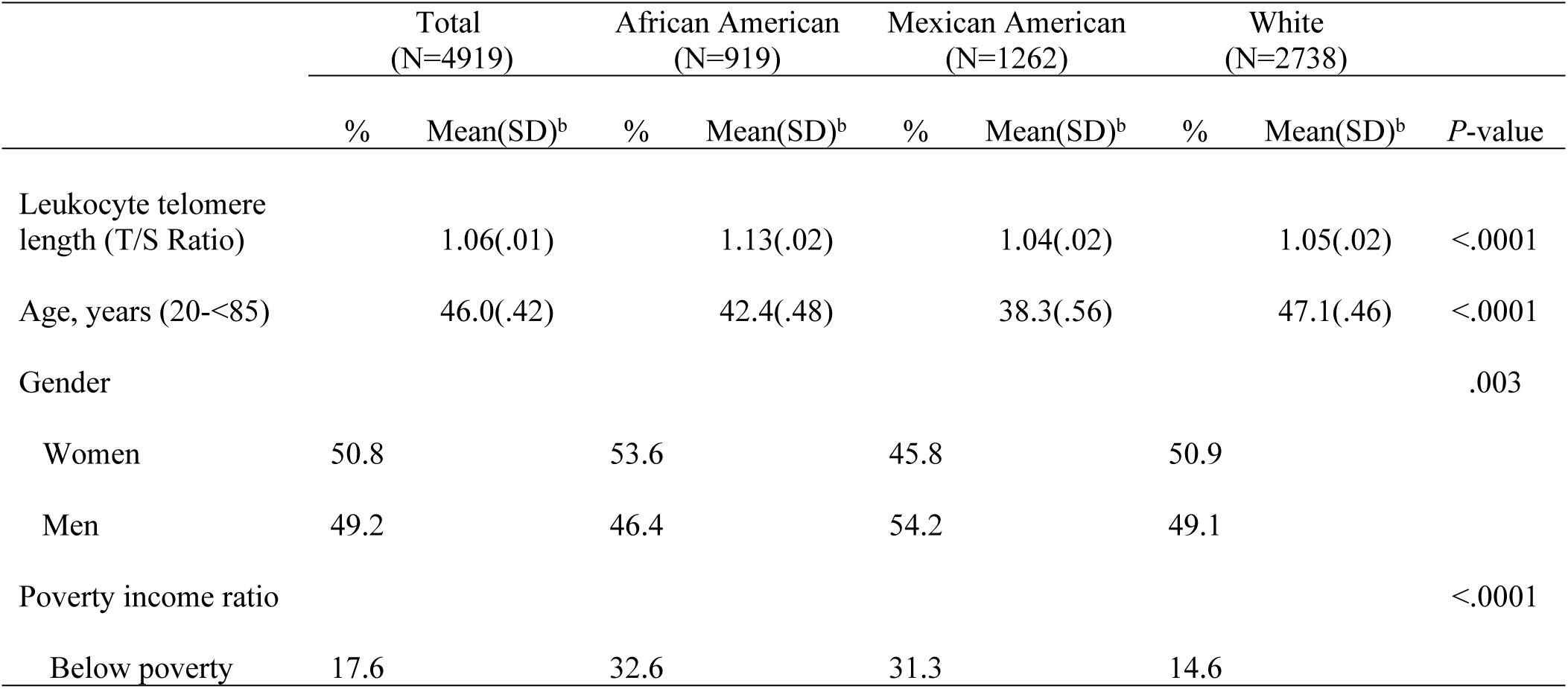
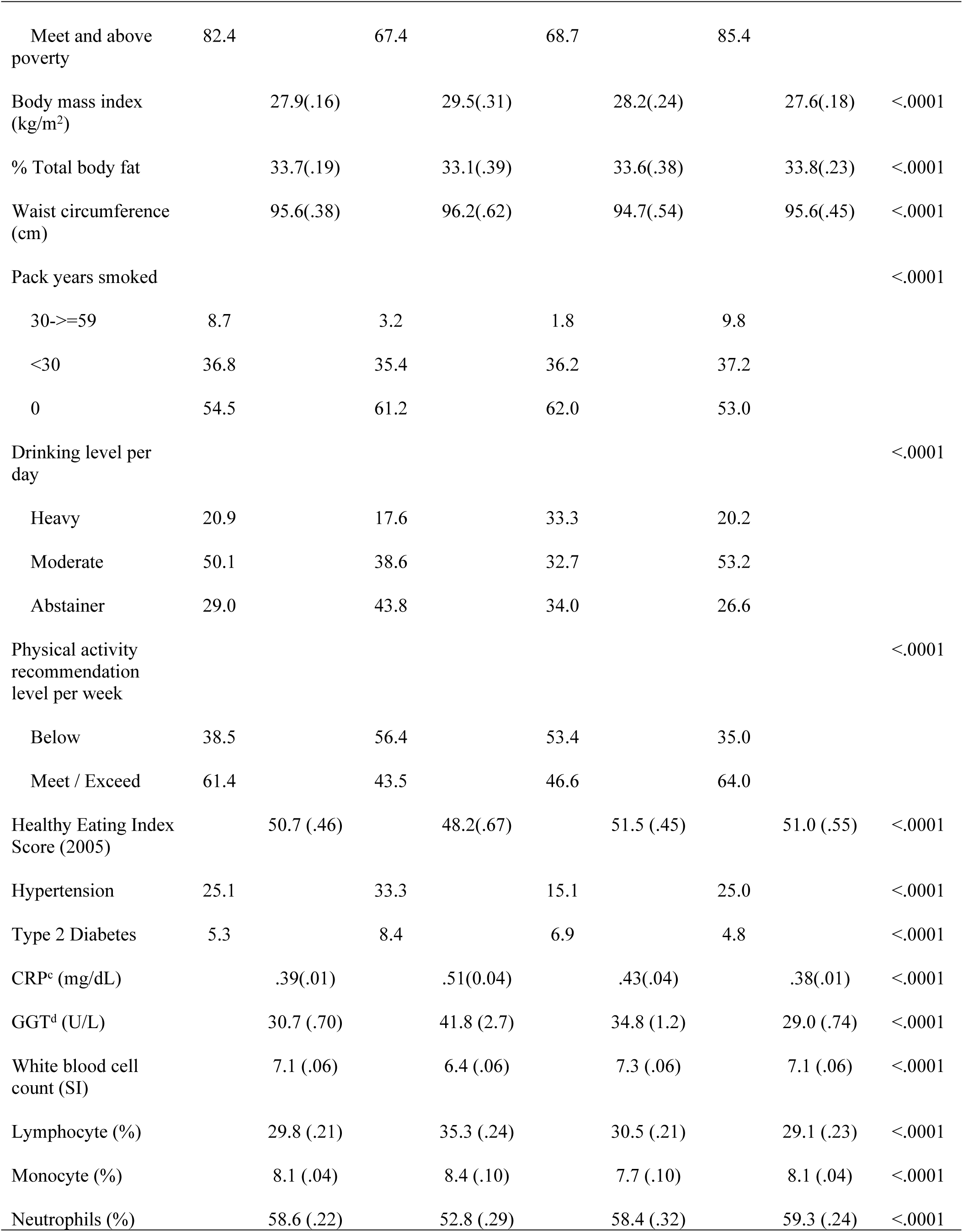
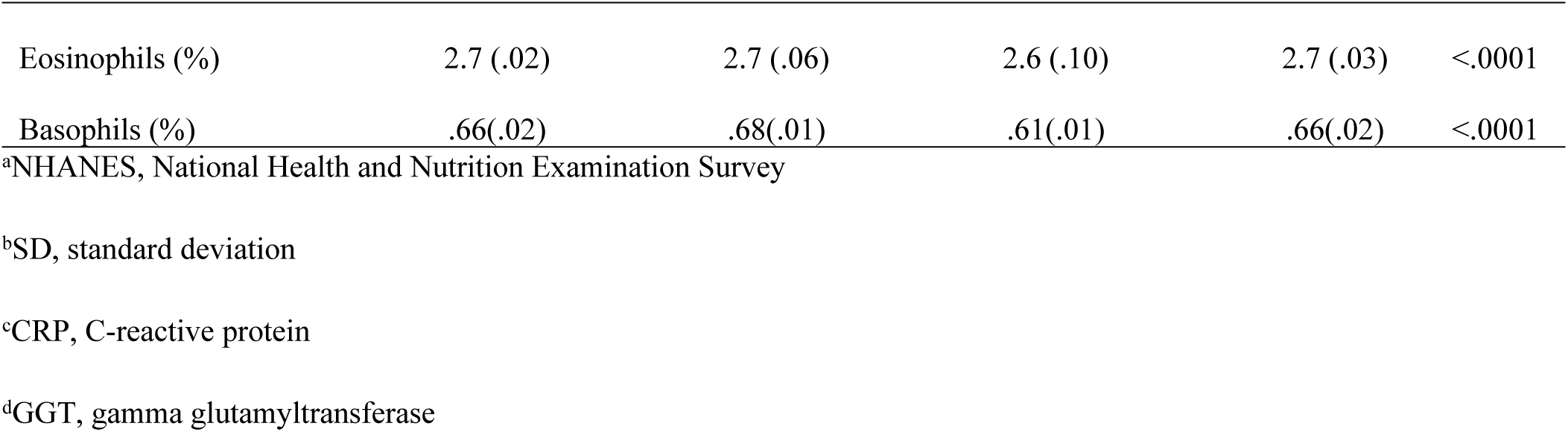
Total and race/ethnic specific weighted characteristics of study variables, NHANES^a^ 1999-2002 (N=4919)

### Adiposity and LTL according to each race/ethnic group

The results for the adjusted association comparing adiposity measures and LTL differences according to each race/ethnic group are presented in Table 2. Findings reveal LTL significantly decreased 28% and 11% for each unit increase in BMI and waist circumference in African Americans, respectively. LTL significantly decreased 33% in Mexican Americans due to increasing % total body fat. Whites experienced a significant 19%, 23%, and .08% decrease in LTL associated with increasing BMI, % total body fat, and waist circumference, respectively. There was no significant association between LTL and % total body fat in African Americans or BMI and waist circumference in Mexican Americans.

**Table 2:**
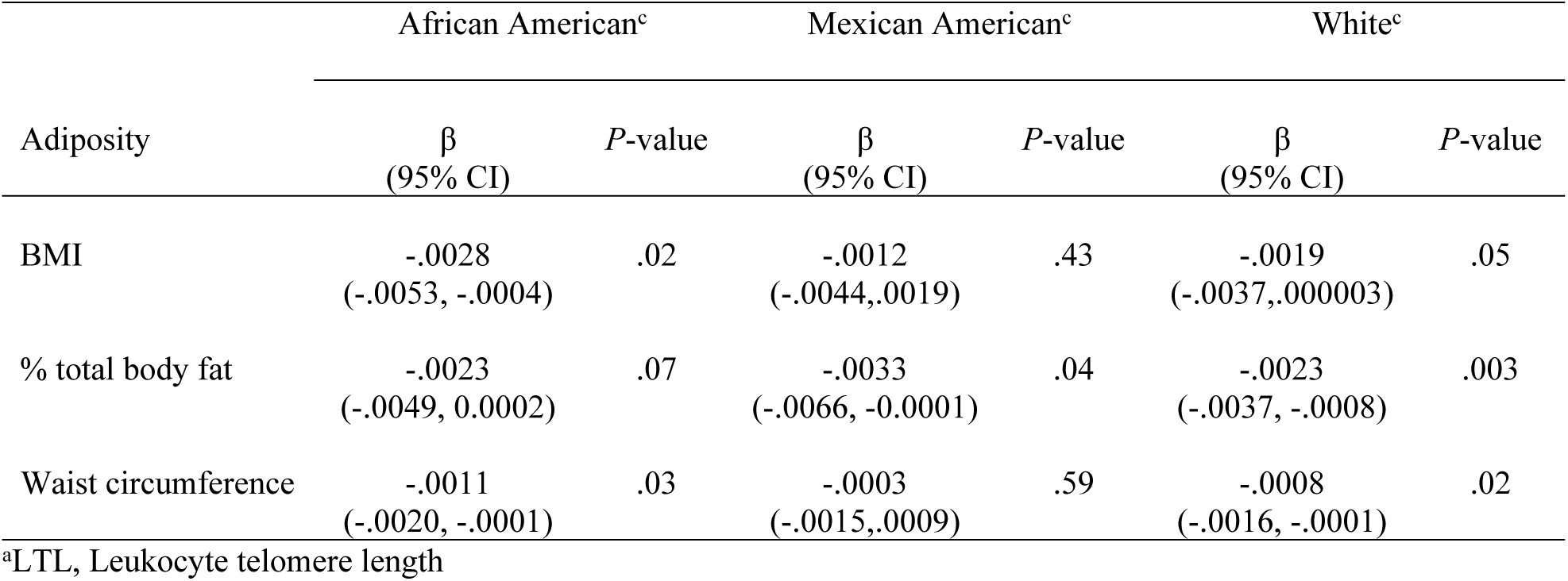
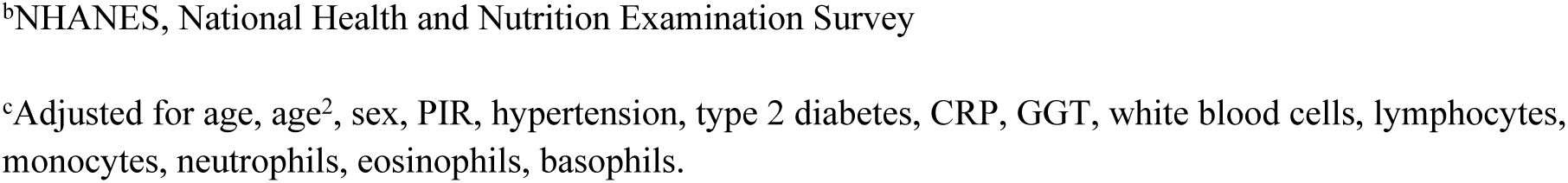
Adjusted ordinary least squares regression of log-transformed LTL^a^ (T/S ratio) on adiposity stratified by individual race/ethnic group, NHANES^b^1999-2002

| Adiposity | African American<sup>c</sup> |  | Mexican American<sup>c</sup> |  | White<sup>c</sup> |  |
| --- | --- | --- | --- | --- | --- | --- |
| | $\beta$<br>(95% CI) | *P*-value | $\beta$<br>(95% CI) | *P*-value | $\beta$<br>(95% CI) | *P*-value |
| BMI | -.0028<br>(-.0053, -.0004) | .02 | -.0012<br>(-.0044,.0019) | .43 | -.0019<br>(-.0037,.000003) | .05 |
| % total body fat | -.0023<br>(-.0049, 0.0002) | .07 | -.0033<br>(-.0066, -0.0001) | .04 | -.0023<br>(-.0037, -.0008) | .003 |
| Waist circumference | -.0011<br>(-.0020, -.0001) | .03 | -.0003<br>(-.0015,.0009) | .59 | -.0008<br>(-.0016, -.0001) | .02 |

<sup>a</sup>LTL, Leukocyte telomere length
<sup>b</sup>NHANES, National Health and Nutrition Examination Survey
<sup>c</sup>Adjusted for age, age<sup>2</sup>, sex, PIR, hypertension, type 2 diabetes, CRP, GGT, white blood cells, lymphocytes, monocytes, neutrophils, eosinophils, basophils.

### Adiposity and LTL by sex specific race/ethnicity

Findings comparing differences in the association of LTL and adiposity measures according to sex specific race/ethnic groups are presented in Table 3. There was no association between any of the adiposity measures and LTL in African American and Mexican American men. Only White men experienced a 26% significant decrease in LTL associated with increasing BMI and increasing waist circumference was marginally associated with a 11% decrease in LTL. African American women experienced a significant 41%, 44%, and 16% decrease in LTL due increasing in BMI, % total body fat and waist circumference, respectively. Increasing % total body fat resulted in a significant 29% decrease in LTL in White women. There was no association with any of the adiposity measures in Mexican American women.

**Table 3:** Adjusted ordinary least squares regression of log-transformed LTL^a^ (T/S ratio) on adiposity comparing race/ethnic group stratified by sex, NHANES^b^ 1999-2002

| Adiposity | Men<sup>c</sup> |  |  |  |  |  |
| --- | --- | --- | --- | --- | --- | --- |
|  | African American<sup>c</sup> |  | Mexican American<sup>c</sup> |  | White<sup>c</sup> |  |
| | $\beta$<br>(95% CI) | *P*-value | $\beta$<br>(95% CI) | *P*-value | $\beta$<br>(95% CI) | *P*-value |
| BMI | -.0016<br>(-.0060, .0028) | .46 | -.0025<br>(-.0079, .0029) | .34 | -.0026<br>(-.0053, .0001) | .05 |
| % total body fat | -.0012<br>(-.0055, .0032) | .58 | -.0041<br>(-.0092, .0011) | .11 | -.0018<br>(-.0042, .0005) | .12 |
| Waist circumference | -.0008<br>(-.0025, .0010) | .37 | -.0009<br>(-.0029, .0011) | .36 | -.0011<br>(-.0022, .0001) | .06 |

Women<sup>c</sup>
|  | African American<sup>c</sup> |  | Mexican American<sup>c</sup> |  | White<sup>c</sup> |  |
| --- | --- | --- | --- | --- | --- | --- |
| Adiposity | $\beta$<br>(95% CI) | *P*-value | $\beta$<br>(95% CI) | *P*-value | $\beta$<br>(95% CI) | *P*-value |
| BMI | -.0041<br>(-.0070, -.0012) | .007 | -.0003<br>(-.0037, .0031) | .85 | -.0017<br>(-.0039, .0005) | .12 |
| % total body fat | -.0044<br>(-.0081, -.0007) | .02 | -.0028<br>(-.0069, .0012) | .16 | -.0029<br>(-.0048, -.0009) | .006 |
| Waist circumference | -.0016<br>(-.0031, -.0001) | .04 | .0003<br>(-.0015, .0020) | .77 | -.0007<br>(-.0017, .0004) | .19 |

<sup>a</sup>LTL, leukocyte telomere length
<sup>b</sup>NHANES, National Health and Nutrition Examination Survey
<sup>c</sup>Adjusted for age, age<sup>2</sup>, PIR, hypertension, type 2 diabetes, CRP, GGT, white blood cells, lymphocytes, monocytes, neutrophils, eosinophils, basophils.

### Moderating and mediator effects

The moderating effect of race/ethnicity and sex was not associated with adiposity measures and LTL as evidenced by non-significant *z* scores. The *z* score for BMI is *z*_AfricanAmerican vs White_ = .03, *P* = .97, *z*_African American vs Mexican Americans_ = -.33, *P* = .74, *z*_White vs Mexican American_ = - .37, *P* = .70. The *z* score for % total body fat is *z*_African American vs White_ = .97, *P* = .32, _*z*African American vs Mexican Americans_ = 1.02, *P* = .30, *z*_White vs Mexican American_ = .20, *P* = .83. The *z* score for waist circumference is *z*_African American vs White_ = .22, *P* = .82, *z*_African American vs Mexican Americans_ = -.39, *P* = .69, *z*_White vs Mexican American_ = -.64, *P* = .52. For sex the *z* score for BMI is *z*_men vs women_ = -.51, *P* = .60, for % total body fat *z*_men vs women_ = .14, *P* = .88, and for waist circumference *z*_men vs women_ = -.49, *P* = .61. There was also no significant interaction for race/ethnicity and sex in the aggregate model (data not presented). Therefore, the test of mediation for health behaviors associated with LTL and adiposity measures was performed on the full sample due to the lack of moderating effects. Table 4 presents the combined results associated with separate linear regression analysis without adjustment for mediators, with adjustment for mediators, and separate test of mediation for each health behavior contributing to the relationship between adiposity measure and LTL. Results revealed BMI and waist circumference were associated with relatively similar significant relationships with decreased LTL after adjustment for mediators as indicated by the *p* values in model 1 and model 2. However, the association between LTL and % total body fat disappeared after adjustment for mediators (*p* = .07).

**Table 4:**
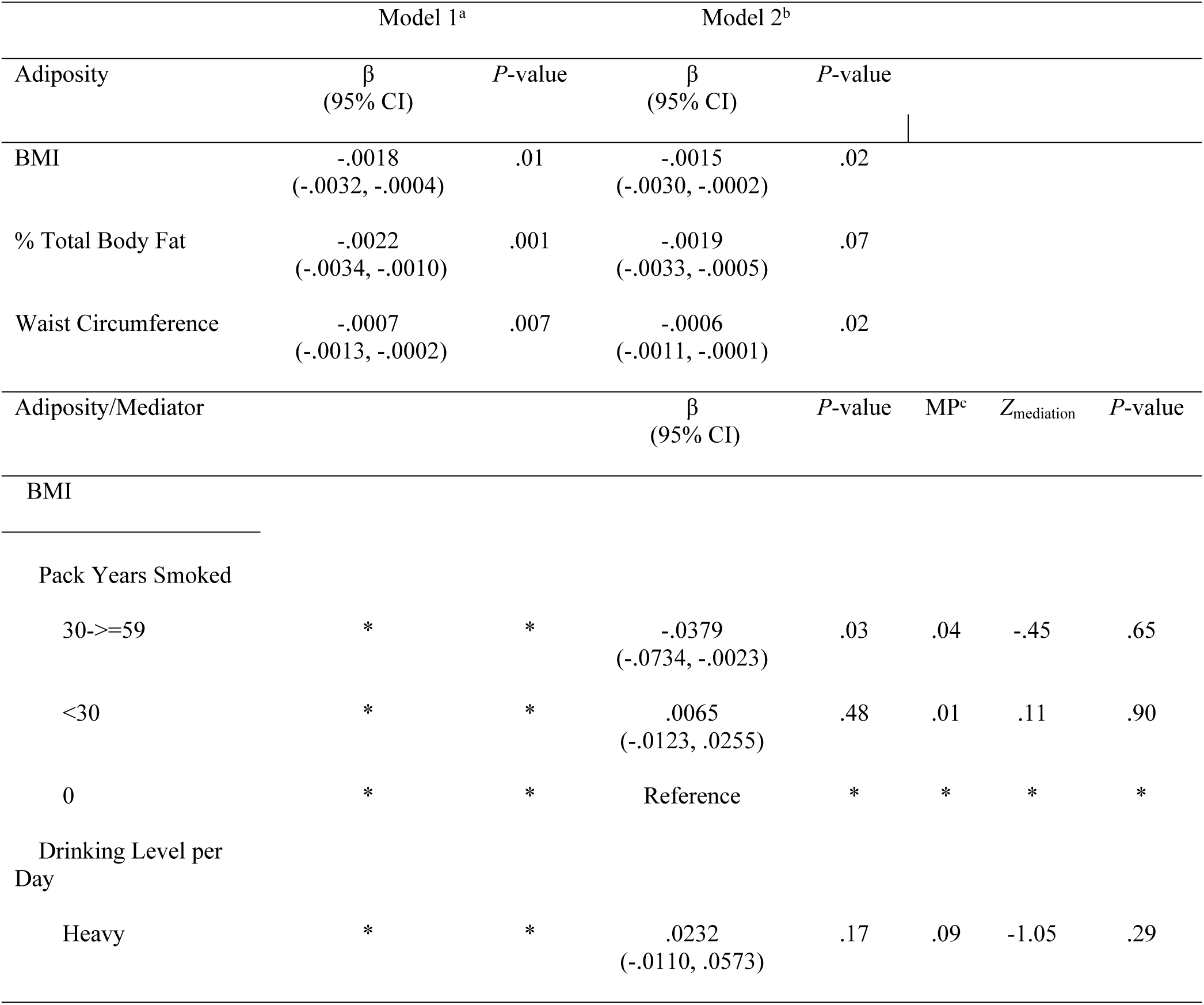
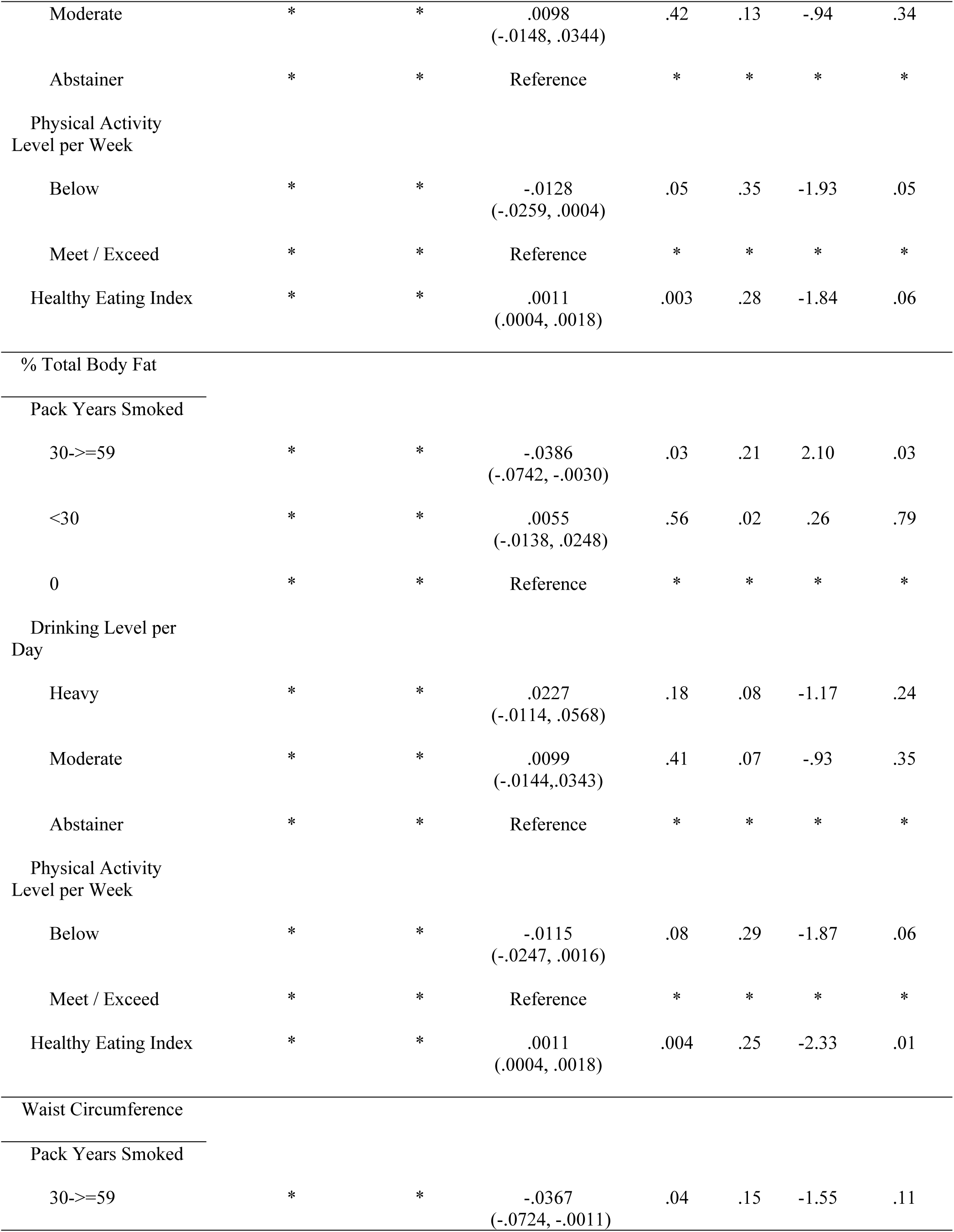
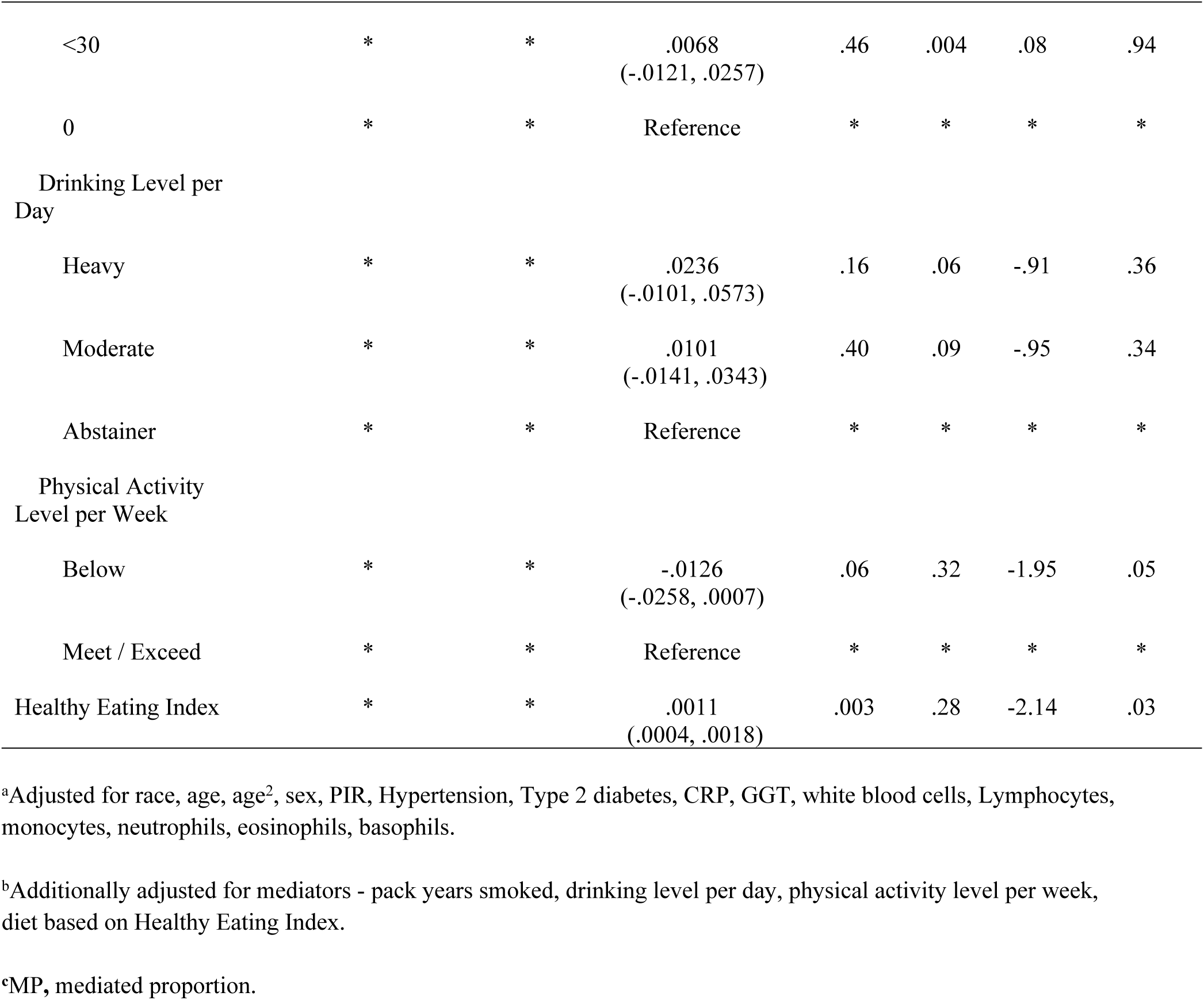
Adjusted ordinary least squares regression of log-transformed LTL (T/S ratio) on adiposity for the total sample without mediators, with mediator effects and the effect of mediators between adiposity and LTL, NHANES 1999

Findings regarding the mediation effects of adverse health behaviors reveal that 30-=>59 pack years smoked was associated with about a 4% decrease in LTL for each unit increase in BMI (*p* = .03); however, it was not a significant mediator (*z*_30-=>59 pack years_ = -.45, *p* = .65). Alcohol consumption per day was also not a significant mediator between LTL and BMI. On the other hand, physical activity below the recommended guidelines was associated with a 1.28% decrease in LTL and BMI (*p* = .05) and accounted for 35% of the relationship (*z*_physical activity_ = −1.92, *p* = .05). Diet as measured by HEI was associated with a 11% increase in LTL for each unit increase in BMI (*p*=.003); however, it was marginally associated with the correlation (*z*_HEI_ = − 1.83, *p* = .06).

Smoking 30-=>59 pack years was associated with a 4% decrease in LTL and increasing % total body fat (*p*=.03) and was a significant mediator contributing 21% of the association (*z*_30-=>59 pack years_ = −2.52, *p* = .03). Physical activity below recommended guideline was marginally correlated with 1.15% decrease in LTL for each unit increase in % total body fat (*p*=.08) and was a marginal mediating factor (*z*_physical activity_ = −1.86, *p* = .06). Diet based HEI resulted in a 11% increase in LTL due to increasing % total body fat (*p*=.004) and was responsible for 25% of the relationship (*z*_HEI_ = −2.32, *p* = .01).

30-=>59 pack years smoked was associated with a 4% decrease in LTL for each unit increase in waist circumference (*p* = .04); however, it was not a significant mechanism between the association (*z*_30-=>59 pack years_ = −1.55, *p*=.11) Physical activity below the recommended guidelines resulted in a 1.26% marginal decrease in LTL and waist circumference (*p*=.06) but was a mediating effect responsible for 32% of the relationship (*z*_physical activity_ = −1.95, *p* = .05). Diet measured by HEI was associated with a 11% increase in LTL and waist circumference (*p*=.003) and significantly contributed 28% to the mechanism between the association (*z*_HEI_ = −2.14, *p* = .03).

## Discussion

The objective of our study was to assess the association between adiposity measures and LTL in a representative US sample population comprised of major racial/ethnic groups and demonstrate mediation effects based on adverse health behaviors. As a group, African Americans experienced shorter telomere length associated with BMI and waist circumference. Whites, on the other hand, experienced shorter telomere length for each of the measures. Only % total body fat correlated with shorter telomere length in Mexican Americans. This finding may reflect how adiposity is concentrated in Mexican Americans. We observed a steeper decline in telomere length associated with BMI in African Americans compared to Whites. It has been demonstrated that regional and whole-body adiposity, including skeletal muscle and bone, differ by race/ethnicity and may not adequately reflect adiposity measures when comparing one race/ethnic group to another.[41] Our stratified findings may support this theory.

When analyzed separately by sex, we found no association with any of the adiposity measures and telomere length in African American or Mexican American men. African American and Mexican American men tend to be leaner compared to African American and Hispanic American women. [42] Only BMI was associated with shorter telomere length in White men. The fact that only one adiposity measure was correlated with shorter telomere length is consistent with this group having a lower overall prevalence of obesity.[42]

We observed opposite findings for women. For instance, African American women experienced shorter telomere length related to increases in each of the adiposity measures. This finding is not surprising given African American women have the highest prevalence of obesity compared to men and women in other racial/ethnic groups.[42] One theory suggests African American women are a unique group susceptible to obesity due to mechanisms associated with dietary preferences and early childbearing.[43] It may also be due to differences in metabolism and perceptions about an ideal body.[44] As with Mexican American men, we found no correlation in Mexican American women associated with any of the exposure measures and telomere length. We were surprise to find the lack of association in Mexican American women given the high prevalence of obesity in Hispanic women.[42, 45] National prevalence rates are generally based on aggregate data of Hispanics in general and does not consider ethnic differences within Hispanics. Our findings on Mexican American women may reflect such heterogeneity and may not be indicative of obesity status in Mexican American women. Only % total body fat was correlated with shorter telomere length in White women. This is not a unique finding given White women have an overall lower prevalence of obesity. [42] The difference in the association by sex that we observed may be due to several factors – including environmental and hormonal.[11, 19]

Our findings regarding adverse health behaviors as mediators revealed physical activity per week below the recommended guideline was a mechanism associated with the relationship between BMI and shorter telomere length. 30-=>59 pack years smoked was a pathway between % total body fat and shorter telomere length while an improvement in diet was associated with an increase in telomere length and % total body fat. Inadequate weekly physical activity was also a causal pathway between waist circumference and shorter telomere. An improvement in diet was a mechanism between longer telomere length and waist circumference. Level of drinking per day was not a mediator between any of the adiposity indices and telomere length.

There are conflicting findings regarding the relationship between adiposity and telomere length in the literature.[26] Lee and colleagues, for instance, studied 345 White individuals in the greater Dayton, Ohio and demonstrated individuals with higher total and abdominal adiposity have shorter telomere length.[12] Findings from two studies using national NHANES data similarly revealed an increase in BMI, waist circumference and % total body fat was associated with a decrease in LTL in the aggregate sample which reflect our findings.[14, 15] An investigation of the Cardiovascular Health Study, on the other hand, found no association between telomere length and BMI and waist circumference.[16] A study by Maceneay et al of 67 middle-aged and older adults also revealed no correlation between normal and overweight/obese BMI parameters with telomere length.[18] A study of 322 postmenopausal women residing in Seattle, Washington revealed no association with BMI and % body fat.[19]

Few studies have compared the association between adiposity and telomere length according to race/ethnicity and concomitant sex. We could find only one study of 317 White and African American adults residing in South Carolina.[17] The investigators found no relationship between BMI and visceral fat in Whites, African Americans or by sex.

Other studies with a racial/ethnic homogenous sample population also produced mixed results. A Swedish study show selective adiposity measures were associated with a decrease in telomere length only in women.[46] An investigation in Denmark revealed an inverse association between BMI and telomere length.[47] A recent study found obesity was related to shorter telomere length in Latina women.[48] However, a study sample of Koreans showed waist circumference was negatively associated with telomere length.[49] These conflicting findings between adiposity and cellular aging as measured by telomere length may be due to several factors-including study design, participant characteristics, a small study sample and other limitations.

Obesity is a major risk factor for chronic morbidities and is due, in large part, to adverse modifiable lifestyle factors.[5] The predominant mechanism through which obesity may shorten telomere length and increase risk of aging-related diseases include increased oxidative stress, which increase telomere erosion, inflammation and accelerates leukocyte turnover.[9-11] A non-biological mechanism contributing to telomere attrition may also include adverse behaviors factors.[26] Several studies have established a parallel relationship between adverse lifestyle factors and telomere length. Patel el al, for instance, found lack of physical activity resulted in shorter telomere length among US adults.[24] A national sample of US women likewise found smoking, unhealthy diet and lower physical activity was correlated with telomere attrition.[25] Other investigations fail to establish a correlation.[19, 21, 22] None have assessed the relationship associated with adverse lifestyle between obesity and telomere length. Our investigation was designed to examine health behaviors as mediators between adiposity and telomere length. Findings demonstrate selective lifestyle behavior factors as a potential causal pathway. Such a relationship suggest improvements in lifestyle may reduce biological aging and prevent telomere cell senescence due to obesity.

## Limitations

There are some caveats to our study that require consideration. First, we do not know the direction of the relationship between adiposity and LTL. Some researchers argue that selective adoption may be a causal factor related to telomere length.[50] Selective adoption could occur either because telomere length directly affects behavior or because behavior affects telomere length, or both are affected by a third variable – such as exposure to early-life adversity. In addition, telomere senescence occurs overtime and may present in some cases with a U-shaped pattern.[51] The NHANES is a cross-sectional survey and changes in LTL may come before exposure. Therefore, the correlations we observed should not be interpreted as causal. One way to address these important issues is to design longitudinal analysis to measure the bi-directional effect of differences in LTL and adiposity overtime before obesity event. Second, although we adjusted for potential confounders – other unmeasured confounding factors may exist resulting in “omitted variable bias” such as heritability, ancestry, menopausal status, adiposity biomarkers (i.e. leptin and adiponectin) and sex-hormones that may affect our findings. Third, we measured telomere length only in leukocytes. Whether our findings can be extrapolated to other tissues is unclear. However, studies have demonstrated robust correlations between LTL and telomere length in other tissues.[52, 53] Fourth, our mediation measures are subject to measurement error even though they have been validated and proven to be accurate in other studies.[54-56]

Despite, these limitations, our study has many strengths. It is comprised of a representative major racial/ethnic sample of US adults from which findings can be extrapolated. It is among the largest and first study to investigate the association of adiposity and telomere length according to race/ethnicity and sex specific race/ethnicity. Finally, we investigated the causal pathway between adiposity and telomere length based on potential modifiable lifestyle behavioral factors. Our detailed measurements of lifestyle and dietary factors enabled us to make categories that were consistent with current guideline on lifestyle and diet which can subsequently be translated into public health intervention messages.

## Conclusion

Telomere length is a measure of biological and cellular aging.[11] Obesity is increasing at an epidemic rate and is associated with several age-related health conditions.[5, 57] Several studies have investigated the relationship between adiposity and telomere length.[12-19] Ours is the first to assess such a relationship in a US representative sample according to race/ethnicity and corresponding sex. Our findings reveal that African Americans and Whites have a worse overall profile regarding the association between collective adiposity measures and telomere length. White men experienced decreased telomere length due to increasing BMI and there was no relationship observed in African American and Mexican American men. White women only experienced shorter telomere length due to increasing % total body fat. African American women experienced shorter telomere length associated with increases in each of the adiposity measures and have a more deleterious health profile based on sex. Our findings also reveal selective adverse lifestyle factors as a mechanism underlying the relationship between adiposity and LTL which portend modifying such factors may result in improvements in cellular and biological aging due to obesity among US adults.

## Author Contributions

### Conceptualization, design, acquisition of data

Sharon K. Davis

### Statistical analysis

Ruihua Xu and Sharon K. Davis

### Interpretation of data

Sharon K. Davis, Ruihua Xu, Rumana J. Khan, Amadou Gaye, Yie Liu

### Writing first draft

Sharon K. Davis

### Writing, review and editing

Sharon K. Davis, Ruihua Xu, Rumana J. Khan, Amadou Gaye, Yie Liu

All authors approved the final version of the manuscript to be published and all agree to be accountable for all aspects of the work in ensuring that questions related to the accuracy or integrity of any part of the work are appropriately investigated and resolved.

## References

1. Sun X, Du T: Trends in cardiovascular risk factors among U.S. men and women with and without diabetes, 1988–2014. BMC Public Health 2017, 17(1):893.

2. Ng M, Fleming T, Robinson M, Thomson B, Graetz N, Margono C, Mullany EC, Biryukov S, Abbafati C, Abera SF et al: Global, regional and national prevalence of overweight and obesity in children and adults 1980-2013: A systematic analysis. Lancet (London, England) 2014, 384(9945):766–781.

3. Available from: http://www.who.int/newd-room/fact-sheets/detail/obesity-and-overweight.

4. Ogden CL, Carroll MD, Kit BK, Km F. Prevalence of obesity among adults: United States, 2011-2012. NCHS data brief, no 131 Hyattsville, MD, National Center for Health Statistics 2013.

5. Poirier P, Giles TD, Bray GA, Hong Y, Stern JS, Pi-Sunyer FX, Eckel RH: Obesity and cardiovascular disease: pathophysiology, evaluation, and effect of weight loss: an update of the 1997 American Heart Association Scientific Statement on Obesity and Heart Disease from the Obesity Committee of the Council on Nutrition, Physical Activity, and Metabolism. Circulation 2006, 113(6):898–918.

6. Hammadah M, Al Mheid I, Wilmot K, Ramadan R, Abdelhadi N, Alkhoder A, Obideen M, Pimple PM, Levantsevych O, Kelli HM et al: Telomere Shortening, Regenerative Capacity, and Cardiovascular Outcomes. Circulation research 2017, 120(7):1130–1138.

7. Fyhrquist F, Saijonmaa O, Strandberg T: The roles of senescence and telomere shortening in cardiovascular disease. Nature reviews Cardiology 2013, 10(5):274–283.

8. Haycock PC, Heydon EE, Kaptoge S, Butterworth AS, Thompson A, Willeit P: Leucocyte telomere length and risk of cardiovascular disease: systematic review and meta-analysis. BMJ (Clinical research ed) 2014, 349:g4227.

9. Suzuki K, Ito Y, Ochiai J, Kusuhara Y, Hashimoto S, Tokudome S, Kojima M, Wakai K, Toyoshima H, Tamakoshi K et al: Relationship between obesity and serum markers of oxidative stress and inflammation in Japanese. Asian Pacific journal of cancer prevention: APJCP 2003, 4(3):259–266.

10. Wolkowitz OM, Mellon SH, Epel ES, Lin J, Dhabhar FS, Su Y, Reus VI, Rosser R, Burke HM, Kupferman E et al: Leukocyte telomere length in major depression: correlations with chronicity, inflammation and oxidative stress--preliminary findings. PloS ONE 2011, 6(3):e17837.

11. Aviv A: Telomeres and human aging: facts and fibs. Science of aging knowledge environment: SAGE KE 2004, 2004(51):pe43.

12. Lee M, Martin H, Firpo MA, Demerath EW: Inverse association between adiposity and telomere length: The Fels Longitudinal Study. American journal of human biology: the official journal of the Human Biology Council 2011, 23(1):100–106.

13. Njajou OT, Cawthon RM, Blackburn EH, Harris TB, Li R, Sanders JL, Newman AB, Nalls M, Cummings SR, Hsueh WC: Shorter telomeres are associated with obesity and weight gain in the elderly. International journal of obesity (2005) 2012, 36(9):1176–1179.

14. Rehkopf DH, Needham BL, Lin J, Blackburn EH, Zota AR, Wojcicki JM, Epel ES: Leukocyte Telomere Length in Relation to 17 Biomarkers of Cardiovascular Disease Risk: A Cross-Sectional Study of US Adults. PLoS Medicine 2016, 13(11):e1002188.

15. Batsis JA, Mackenzie TA, Vasquez E, Germain CM, Emeny RT, Rippberger P, Lopez-Jimenez F, Bartels SJ: Association of adiposity, telomere length and mortality: data from the NHANES 1999-2002. International journal of obesity (2005) 2018, 42(2):198–204.

16. Fitzpatrick AL, Kronmal RA, Gardner JP, Psaty BM, Jenny NS, Tracy RP, Walston J, Kimura M, Aviv A: Leukocyte telomere length and cardiovascular disease in the cardiovascular health study. American journal of epidemiology 2007, 165(1):14–21.

17. Diaz VA, Mainous AG, Player MS, Everett CJ: Telomere length and adiposity in a racially diverse sample. International journal of obesity (2005) 2010, 34(2):261–265.

18. MacEneaney OJ, Kushner EJ, Westby CM, Cech JN, Greiner JJ, Stauffer BL, DeSouza CA: Endothelial progenitor cell function, apoptosis, and telomere length in overweight/obese humans. Obesity (Silver Spring) 2010, 18(9):1677–1682.

19. Mason C, Risques R-A, Xiao L, Duggan CR, Imayama I, Campbell KL, Kong A, Foster-Schubert KE, Wang CY, Alfano CM et al: Independent and Combined Effects of Dietary Weight Loss and Exercise on Leukocyte Telomere Length in Postmenopausal Women. Obesity (Silver Spring, Md) 2013, 21(12):E549–E554.

20. Mundstock E, Sarria EE, Zatti H, Mattos Louzada F, Kich Grun L, Herbert Jones M, Guma FT, Mazzola In Memoriam J, Epifanio M, Stein RT et al: Effect of obesity on telomere length: Systematic review and meta-analysis. Obesity (Silver Spring) 2015, 23(11):2165–2174.

21. Rode L, Bojesen SE, Weischer M, Nordestgaard BG: High tobacco consumption is causally associated with increased all-cause mortality in a general population sample of 55,568 individuals, but not with short telomeres: a Mendelian randomization study. International journal of epidemiology 2014, 43(5):1473–1483.

22. Weischer M, Bojesen SE, Nordestgaard BG: Telomere shortening unrelated to smoking, body weight, physical activity, and alcohol intake: 4,576 general population individuals with repeat measurements 10 years apart. PLoS genetics 2014, 10(3):e1004191.

23. Zhang C, Lauderdale DS, Pierce BL: Sex-Specific and Time-Varying Associations Between Cigarette Smoking and Telomere Length Among Older Adults. American journal of epidemiology 2016, 184(12):922–932.

24. Patel CJ, Manrai AK, Corona E, Kohane IS: Systematic correlation of environmental exposure and physiological and self-reported behaviour factors with leukocyte telomere length. International journal of epidemiology 2017, 46(1):44–56.

25. Sun Q, Shi L, Prescott J, Chiuve SE, Hu FB, De Vivo I, Stampfer MJ, Franks PW, Manson JE, Rexrode KM: Healthy lifestyle and leukocyte telomere length in U.S. women. PloS one 2012, 7(5):e38374.

26. Mundstock E, Zatti H, Louzada FM, Oliveira SG, Guma FT, Paris MM, Rueda AB, Machado DG, Stein RT, Jones MH et al: Effects of physical activity in telomere length: Systematic review and meta-analysis. Ageing research reviews 2015, 22:72–80.

27. Ford ES, Zhao G, Tsai J, Li C: Low-risk lifestyle behaviors and all-cause mortality: findings from the National Health and Nutrition Examination Survey III Mortality Study. American journal of public health 2011, 101(10):1922–1929.

28. Aviv A, Valdes AM, Spector TD: Human telomere biology: pitfalls of moving from the laboratory to epidemiology. International journal of epidemiology 2006, 35(6):1424–1429.

29. Cawthon RM: Telomere measurement by quantitative PCR. Nucleic acids research 2002, 30(10):e47.

30. Gunter EW LB, Koncikowski SM: Laboratory procedures used for the Third National Health and Nutrition Examination Survey (NHANES III), 1988-1994. In. Edited by US Department of Health and Human Services NCfHS; 1996.

31. Mannino DM, Aguayo SM, Petty TL, Redd SC: Low lung function and incident lung cancer in the United States: data From the First National Health and Nutrition Examination Survey follow-up. Archives of internal medicine 2003, 163(12):1475–1480.

32. Physical Activity Guidelines for Americans [https://health.gov/paguidelines/guidelines/]

33. Guenther PM, Reedy J, Krebs-Smith SM: Development of the Healthy Eating Index-2005. Journal of the American Dietetic Association 2008, 108(11):1896–1901.

34. US Census Bureau, Population Division, Fertility & Family Statistics Branch. In. Edited by Bureau UC; 2004.

35. National Center for Health Statistics. Analytic and reporting guidelines: The National Health and Nutrition Examination Survey (NHANES). In. Hyattsville, MD: National Center for Health Statistics, Centers for Disease Control and Prevention; 2006.

36. Paternoster R BR, Mazerolle P, Piquero A: Using the correct statistical test for the quality of regression coefficients. Criminology 1998, 36:859–866.

37. Baron RM, Kenny DA: The moderator-mediator variable distinction in social psychological research: conceptual, strategic, and statistical considerations. Journal of personality and social psychology 1986, 51(6):1173–1182.

38. D I: Mediation analysis and categorical variables: The final froniter. Journal of Consumer Psychology 2012, 22:582–594.

39. Ditlevsen S, Keiding N, Christensen U, Damsgaard MT, Lynch J: Mediation proportion. Epidemiology (Cambridge, Mass) 2005, 16(4):592.

40. SAS 9.3. In. Edited by Inc SI. Cary, NC 2011.

41. Heymsfield SB, Peterson CM, Thomas DM, Heo M, Schuna JM, Jr.: Why are there race/ethnic differences in adult body mass index-adiposity relationships? A quantitative critical review. Obesity reviews: an official journal of the International Association for the Study of Obesity 2016, 17(3):262–275.

42. National Center for Health Statistics. Health, United States, 2015: With Special Feature on Racial and Ethnic Health Disparities. In. Hyattsville, MD.; 2016.

43. Seamans MJ, Robinson WR, Thorpe RJ, Jr., Cole SR, LaVeist TA: Exploring racial differences in the obesity gender gap. Annals of epidemiology 2015, 25(6):420–425.

44. Lee J, Sa J, Chaput JP, Seo DC, Samuel T: Racial/ethnic differences in body weight perception among U.S. college students. Journal of American college health: J of ACH 2018:1–9.

45. Patel SR, Blackwell T, Redline S, Ancoli-Israel S, Cauley JA, Hillier TA, Lewis CE, Orwoll ES, Stefanick ML, Taylor BC et al: The association between sleep duration and obesity in older adults. International journal of obesity (2005) 2008, 32(12):1825–1834.

46. Nordfjall K, Eliasson M, Stegmayr B, Melander O, Nilsson P, Roos G: Telomere length is associated with obesity parameters but with a gender difference. Obesity (Silver Spring) 2008, 16(12):2682–2689.

47. Rode L, Nordestgaard BG, Weischer M, Bojesen SE: Increased body mass index, elevated Creactive protein, and short telomere length. The Journal of clinical endocrinology and metabolism 2014, 99(9):E1671–1675.

48. Wojcicki JM, Elwan D, Lin J, Blackburn E, Epel E: Chronic Obesity and Incident Hypertension in Latina Women Are Associated with Accelerated Telomere Length Loss over a 1-Year Period. Metabolic syndrome and related disorders 2018.

49. Kim J-H, Kim HK, Ko J-H, Bang H, Lee D-C: The Relationship between Leukocyte Mitochondrial DNA Copy Number and Telomere Length in Community-Dwelling Elderly Women. PloS one 2013, 8(6):e67227.

50. Bateson M, Nettle D: Why are there associations between telomere length and behaviour? Philosophical transactions of the Royal Society of London Series B, Biological sciences 2018, 373(1741).

51. Raschenberger J, Kollerits B, Titze S, Kottgen A, Barthlein B, Ekici AB, Forer L, Schonherr S, Weissensteiner H, Haun M et al: Do telomeres have a higher plasticity than thought? Results from the German Chronic Kidney Disease (GCKD) study as a high-risk population. Experimental gerontology 2015, 72:162–166.

52. Okuda K, Bardeguez A, Gardner JP, Rodriguez P, Ganesh V, Kimura M, Skurnick J, Awad G, Aviv A: Telomere length in the newborn. Pediatric research 2002, 52(3):377–381.

53. Wilson WR, Herbert KE, Mistry Y, Stevens SE, Patel HR, Hastings RA, Thompson MM, Williams B: Blood leucocyte telomere DNA content predicts vascular telomere DNA content in humans with and without vascular disease. European heart journal 2008, 29(21):2689–2694.

54. Needham BL, Adler N, Gregorich S, Rehkopf D, Lin J, Blackburn EH, Epel ES: Socioeconomic status, health behavior, and leukocyte telomere length in the National Health and Nutrition Examination Survey, 1999-2002. Social science & medicine (1982) 2013, 85:1–8.

55. Shiao SPK, Grayson J, Lie A, Yu CH: Predictors of the Healthy Eating Index and Glycemic Index in Multi-Ethnic Colorectal Cancer Families. Nutrients 2018, 10(6).

56. Kay MC, Carroll DD, Carlson SA, Fulton JE: Awareness and knowledge of the 2008 Physical Activity Guidelines for Americans. Journal of physical activity & health 2014, 11(4):693–698.

57. Collaboration GBDO, Ng M, Fleming T, Robinson M, Thomson B, Graetz N, Margono C, Mullany EC, Biryukov S, Abbafati C et al: Global, regional and national prevalence of overweight and obesity in children and adults 1980-2013: A systematic analysis. Lancet (London, England) 2014, 384(9945):766–781.

